# The RNA-binding landscape of HAX1 protein indicates its involvement in ribosome biogenesis and translation

**DOI:** 10.1101/2022.01.14.476349

**Authors:** Anna Balcerak, Ewelina Macech-Klicka, Maciej Wakula, Rafal Tomecki, Krzysztof Goryca, Malgorzata Rydzanicz, Mateusz Chmielarczyk, Malgorzata Szostakowska-Rodzos, Marta Wisniewska, Filip Lyczek, Aleksandra Helwak, David Tollervey, Grzegorz Kudla, Ewa A. Grzybowska

## Abstract

HAX1 is a human protein with no known homologues or structural domains, mutations in which cause severe congenital neutropenia through mechanisms that are poorly understood. Previous studies reported RNA-binding capacity of HAX1, but the role of this binding in physiology and pathology remains unexplained. Here we report transcriptome-wide characterization of HAX1 RNA targets using RIP-seq and CRAC, indicating that HAX1 binds transcripts involved in ribosome biogenesis and rRNA processing. Using CRISPR knockouts we find that RNA targets of HAX1 partially overlap with transcripts downregulated in *HAX1* KO, implying a role in mRNA stabilization. Gene ontology analysis demonstrated that genes differentially expressed in *HAX1* KO (including genes involved in ribosome biogenesis and translation) are also enriched in a subset of genes whose expression correlates with *HAX1* expression in four analyzed neoplasms. Functional connection to ribosome biogenesis was also demonstrated by gradient sedimentation ribosome profiles, which revealed differences in the small subunit:monosome ratio in *HAX1* WT/KO. We speculate that changes in HAX1 expression may be important for the etiology of HAX1-linked diseases through dysregulation of translation.

## INTRODUCTION

HAX1 is known as an anti-apoptotic protein with a role in the regulation of cell migration, cell adhesion and calcium homeostasis (Fadeel and Grzybowska 2009). HAX1 deficiency, due to the mutations in *HAX1* gene results in the autosomal recessive severe congenital neutropenia (SCN) called Kostmann disease (Klein et al. 2007). This myelopoietic disorder is caused by granulocyte maturation arrest and the resulting paucity of mature neutrophils, which leads to life-threatening infections. This effect has been attributed to the excessive apoptosis caused by HAX1 deficiency, but the exact molecular mechanism was not demonstrated. Conversely, HAX1 overexpression was documented in several neoplasms (Kwiecinska et al. 2011; Li et al. 2015; Trebinska-Stryjewska et al. 2019; Liang et al. 2020). In here, we would like to propose that these effects may be linked to ribosome dysfunction.

To date, RNA-binding propensity of HAX1 was reported in two particular cases: for vimentin transcript (Al-Maghrebi et al. 2002) and DNA polymerase beta transcript (Sarnowska et al. 2007). Both instances pertained to hairpin structures at the 3’UTR of these transcripts, although the two structures were not similar. In our recent report (Wakula et al. 2020), in which we described HAX1 protein interactome, we also suggested the possibility of RNA-binding by HAX1 deduced from neighboring proteins (first observed by Brannan et al.,(Brannan et al. 2016)).

The role of HAX1 RNA-binding in cellular processes has not been clarified so far, except some suggestions of its involvement in the regulation of specific mRNAs (Grzybowska et al. 2013; Zayat et al. 2015).

HAX1 protein has no homologues or known domains, except for a PEST sequence, which indicates susceptibility for degradation (Li et al. 2012). The presence of BCL2-like domains was disproved (Jeyaraju et al. 2009). Accordingly, HAX1 does not possess any known RNA-binding domain, and a large proportion of the protein is predicted to be disordered (Balcerak et al. 2017; Larsen et al. 2020), so RNA–binding probably occurs in a non-conventional manner.

The current study provides for the first time a comprehensive analysis of HAX1 RNA-interactome, with two independent approaches for the isolation of its RNA targets. Subsequent analysis of the impact of HAX1 on cell transcriptome and analysis of expression in several cancer databases produced coherent results indicating unanticipated HAX1 role in ribosome biogenesis and translation.

Comparison of the experimental data obtained for HAX1 RNA-binding and transcriptome profiling indicates that HAX1 may regulate stability of the bound transcripts. These results were corroborated by the observation that *HAX1* KO affects ribosomal profile, especially concerning the ratio of small ribosomal subunit to monosome. HAX1 involvement in ribosome biogenesis and translation emerging from this work may help to elucidate its many-sided effects on cellular processes and HAX1-associated diseases.

## MATERIAL AND METHODS

### 1. Generation of cell lines

#### 1.1. HEK293FlpInTRex with induced *HAX1* overexpression

##### Plasmid design and molecular subcloning

*HAX1* CDS was obtained by PCR and cloned into prepared vectors with special gene coding tag (Protein A fragment, TEV protease cleavage site and his-tag) in two orientations (tag on 3’ or 5’ end of the *HAX1* coding sequence).

##### Cell line generation

HEK293-FlpIn cells (ThermoFisher Scientific) were transfected with empty vector (negative control, NC) or plasmid with *HAX1* gene with tag on 3’end or 5’end of the gene (LipofectamineTM2000, ThermoFisher Scientific). Cells were detached and seeded on 100mm plates to grow single colonies (selection: Blasticidin 15 µg/ml and Higromycin B 100 µg/ml) for 2 weeks. Single colonies were passaged to 24-well plates and tested by Western blot and qPCR with and without doxycycline induction (18-48 h).

#### 1.2. HL-60 *HAX1* CRISPR knock-out

##### Plasmid design and molecular subcloning

two pairs (4 oligonucleotides) of small guide RNA (sgRNA) complementary to *HAX1* gene were designed using online bioinformatic tool (https://CRISPR.mit.edu). Each of sgRNA pair was introduced to AIO-GFP plasmid encoding EGFP-tagged Cas9 nickase (D10A), as described in (Chiang et al. 2016). Briefly, each of 4 oligonucleotides contained 5’ overhangs (forward: ACCG, reverse: AAAC) compatible with BbsI and BsaI restriction enzymes. BbsI site present in AIO-GFP was utilized to introduce anti-sense (LC - left CRISPER) oligonucleotides and therefore generate AIO-GFP HAX1 LC1 and AIO-GFP HAX1 LC2 plasmids. To each respective plasmid, second (sense, RC - right CRISPER) sgRNA of given pair was introduced utilizing BsaI site. As result two different constructs were generated, AIO-GFP HAX1 LCRC1 and AIO-GFP HAX1 LCRC2.

##### Cell line generation

HL-60 cell line was grown in RPMI1640 medium with L-glutamine (Biowest) and 10% Fetal Bovine Serum (Gibco) at 37°C in a 5% CO_2_. Electroporation of 5×10^6^ HL-60 cells was performed with AIO-GFP HAX1 LCRC1 and AIO-GFP HAX1 LCRC2 using CLB-Transfection™ Kit (Lonza, Austria) and CLB-Transfection™ System (Lonza, Austria) with default program 9 setting. Single transfected cells were sorted to separate wells of 96-well culture plate using BD FACSAria™ III (Becton Dickinson, USA). Cells were cultured in RPMI for 14-21 days, colonies were propagated and successful KO was validated by Western blot in four cell lines, two of which were used in experiments (Figure S1).

### 2. RIP-seq

HL-60 promyelocytic cell line (DSMZ, Germany) was grown in RPMI1640 medium with L-glutamine (Biowest) and 10% Fetal Bovine Serum (Gibco) at 37°C in a 5% CO_2_. The experiment was conducted with EZ-Magna RIP RNA-Binding Protein Immunoprecipitation Kit (Millipore,17-700 Sigma-Aldrich) according to the manufacturer’s protocol. A single freeze-thaw was employed, to gently lyse the cells as described by Keene et al, (Keene et al. 2006). 30×10^6^ cells per sample were collected by centrifugation at 966 x g, 5 minutes at 4°C and washed two times with PBS containing protease inhibitors, resuspended in 200 μL of RIP lysis buffer (Millipore) containing protease and RNase inhibitors, incubated for 5 minutes on ice, snap-frozen in liquid nitrogen and stored at -80°C. Magnetic Beads Protein A/G (from Magna RIP Kit) were incubated overnight (4°C) with 10 µg of anti-HAX1 rabbit polyclonal antibody (Thermo Fisher Scientific, MA, U.S.A.; cat. PA5-27592) and Rabbit IgG (Millipore) as a negative control. The lysates from the previous step were quickly thawed, centrifuged at 14,000 rpm for 10 minutes at 4°C and 150 μL of supernatants were added to each antibody complex in RIP Immunoprecipitation Buffer. The final volume of the immunoprecipitation reaction was 1.5 mL. 10% of the sample was taken and stored as total input. The lysate was incubated with antibody-coated beads for 4 hours at 4°C. After immunoprecipitation, the beads were washed 5 times with 1 mL of cold RIP Wash Buffer. The last, sixth wash was performed with 0.5 mL of wash buffer and 50 μL out of 500 μL of each beads’ suspension was taken to test the efficiency of immunoprecipitation by Western blotting. The remaining 450 μL of each suspension was collected with magnetic separator, immune-complexes and input were eluted and treated with proteinase K (55°C for 30 minutes). RNA was purified with phenol/chloroform extraction followed by ethanol precipitation.

### 3. Sample preparation (RIPseq, RNAseq)

RIP-seq. Concentration of the precipitated RNA samples were checked using QuantiFluor RNA System (Promega) and the samples were used for library preparation. RNA-seq. HL-60 cell lines (WT and *HAX1* KO#1 and #2) were grown to 3.5 × 10^6^ cells in each culture (each cell line in four replicates). RNA was isolated using RNA PureLink Mini (Thermofisher, USA). Genomic DNA was removed from samples using TURBO DNA-free kit (Thermofisher, USA). RNA integrity was assessed using Agilent RNA 6000 Nano Kit (Agilent Technologies, USA). RNA samples with RIN score ≥ 9 were used for the preparation of cDNA libraries.

### 4. NGS library preparation and sequencing (RIP-seq, RNA-seq)

cDNA libraries were prepared using TruSeq™ Stranded Total RNA Library Prep Gold (Illumina, USA) according to manufacturer’s procedure. The average size of the libraries was determined utilizing Agilent 2100 Bioanalyzer and High Sensitivity DNA Kit (Agilent Technologies, USA), while concentration was assessed using Qubit Fluorometer and dsDNA HS Assay Kit (Thermo Fisher Scientific, USA). Uniquely indexed libraries were pooled, mixed with Illumina PhiX Control v3 Library (1% of total amount), and sequenced on HiSeq 1500 (Illumina) on Rapid Run Mode. Single-read sequencing (1x 50 bp) and paired-end sequencing (2×100 bp) was performed for RIP-seq and RNA-seq, respectively.

### 5. NGS data analysis (RIP-seq, RNA-seq)

Raw sequences were trimmed according to quality using Trimmomatic (Bolger et al. 2014) (version 0.39) using default parameters, except MINLEN, which was set to 50. Trimmed sequences were mapped to the human reference genome provided by ENSEMBL, (version grch38_snp_tran) using Hisat2 (Kim et al. 2015) with default parameters. Optical duplicates were removed using MarkDuplicates tool from GATK (McKenna et al. 2010) package (version 4.1.2.0) with default parameters except OPTICAL_DUPLICATE_PIXEL_DISTANCE set to 12000. Mapped reads were associated with transcripts from grch38 database (Zerbino et al. 2018) (Ensembl, version 96) using HTSeq-count (Anders et al. 2015) (version 0.9.1) with default parameters. except –stranded set to “reverse”. Differentially expressed genes were selected using DESeq2 package (Love et al. 2014) (version 1.16.1). Fold change was corrected using apeglm.

### 6. CRAC (Crosslinking and Analyses of cDNAs)

#### Sample preparation

Established stable cell lines with HAX1 overexpression (coding protein tagged on C or N terminal end) were induced with 1 μg/ml doxycycline and incubated 18-48 h. After incubations cells were UV-crosslinked in Stratalinker 1800 (E=400 mJ/cm2) at wavelenght 254 nm. Cells were lysed for 10 min on ice in lysis buffer (50 mM Tris-HCl pH 7.5, 300 mM NaCl, 1% NP-40, 5mM EDTA, 10% glycerol, 5 mM β-mercaptoethanol). Lysates were spun at 4600 RPM, 4ºC for 5 minutes and subsequently filtered using syringe filter 0.45 µm with PES membrane. IgG Sepharose was added to lysates and incubated overnight at 4ºC with rotation. Beads were washed with IgG wash buffer (50 mM Tris-HCl pH 7.5, 800 mM NaCl, 0.5% NP-40, 5 mM MgCl_2_) and PNK wash buffer (25 mM Tris-HCl pH 7.5, 50 mM NaCl, 0.1% NP-40, 1 mM MgCl_2_). RNAs were trimmed on beads using 1 unit of RNAce-IT in 0.45 ml PNK buffer for 7 minutes at 37°C. To stop the reaction supernatant with RNaceIT was removed and the beads were re-suspended in the room temperature denaturing elution buffer Ni-WBI (50 mM Tris-HCl pH 7.5, 300 mM NaCl, 1.5 mM MgCl_2_, 10 mM Imidazole pH 8.0, 0.1 % NP-40 and 6M guanidine hydrochloride). Elution was repeated one more time, both fractions combined and Ni-NTA beads were added for overnight incubation at 4ºC. NI-NTA beads were washed and transferred to Pierce columns. RNA was dephosphorylated with 8U of Thermosensitive Alkaline Phosphatase (Promega) in supplied MultiCore buffer with 80 U RNAsin for 30 minutes at 37ºC. Beads were washed with Ni-WBI and PNK wash buffer. 3’ linker ligation was performed overnight at 16°C with 1 μM 3’linker, 800U of truncated T4 RNA ligase 2 K227Q (New England Biolabs) in supplied PNK buffer with RNAsin (Promega) and 10% PEG8000). Beads were washed with WBI and PNK wash buffer. RNA-protein complexes were radioactively labeled with 32P-γ-ATP (20 μCi) using 40U T4 Polynucleotide Kinase (New England Biolabs) in supplied PNK buffer for 30 min at 37ºC. 5’ linker ligation was performed in the same reaction mixture by addition of 5’ linker to the final 2.5 mM, non-radioactive ATP to final 1.25 mM and 40U T4 RNA ligase 1 for 8 hours at 16ºC. After washing beads with Ni-WBI and PNK wash buffer, RNA-protein complexes were eluted with elution buffer (50 mM Tris-Hcl pH 7.8, 300 mM NaCl, 150 mM Imidazole pH 8.0, 0.1 % NP-40, 5 mM 2-mercaptoethanol) at room temperature for 5 minutes. Protein-RNA complexes were precipitated with 80% acetone in the presence of GlycoBlue at -20°C overnight and spun for 20 minutes with max speed at 4°C. Pellets were resuspended in LDS sample buffer (ThermoFisher), DTT and EDTA and denatured for 3 minutes at 90°C.

#### Autoradiography

Samples were resolved on 4-12% Bis-Tris NuPAGE gel at constant voltage (120V) using NuPAGE MOPS SDS running buffer (Thermo Fisher Scientific) and transferred to nitrocellulose membrane in NuPAGE transfer buffer using BioRad Protean wet-transfer system at constant voltage (100V) for 1 h. Exposition was performed at -80ºC overnight.

#### RNA isolation from membrane

Bands corresponding to RNA crosslinked to the HAX1 protein (Figure 1D) were cut out and incubated with 450 μl Proteinase K buffer (50 mM Tris-HCl pH 7.8, 50 mM NaCl,10 mM imidazole pH 8.0, 0.1% NP-40, 1% SDS, 5 mM EDTA, 5 mM 2-mercaptoethanol) and 200 μg Proteinase K for 2 h at 55ºC. 3M sodium acetate pH 5,2 was added to final 10% and RNA was extracted with 500 µl phenol:chloroform:isoamyl alcohol. After 5 minute spin with maximum speed at 4°C, aqueous phase was collected to a new tube and RNA was precipitated with 3 volumes of Ethanol in the presence of GlucoBlue.

**Figure 1.**
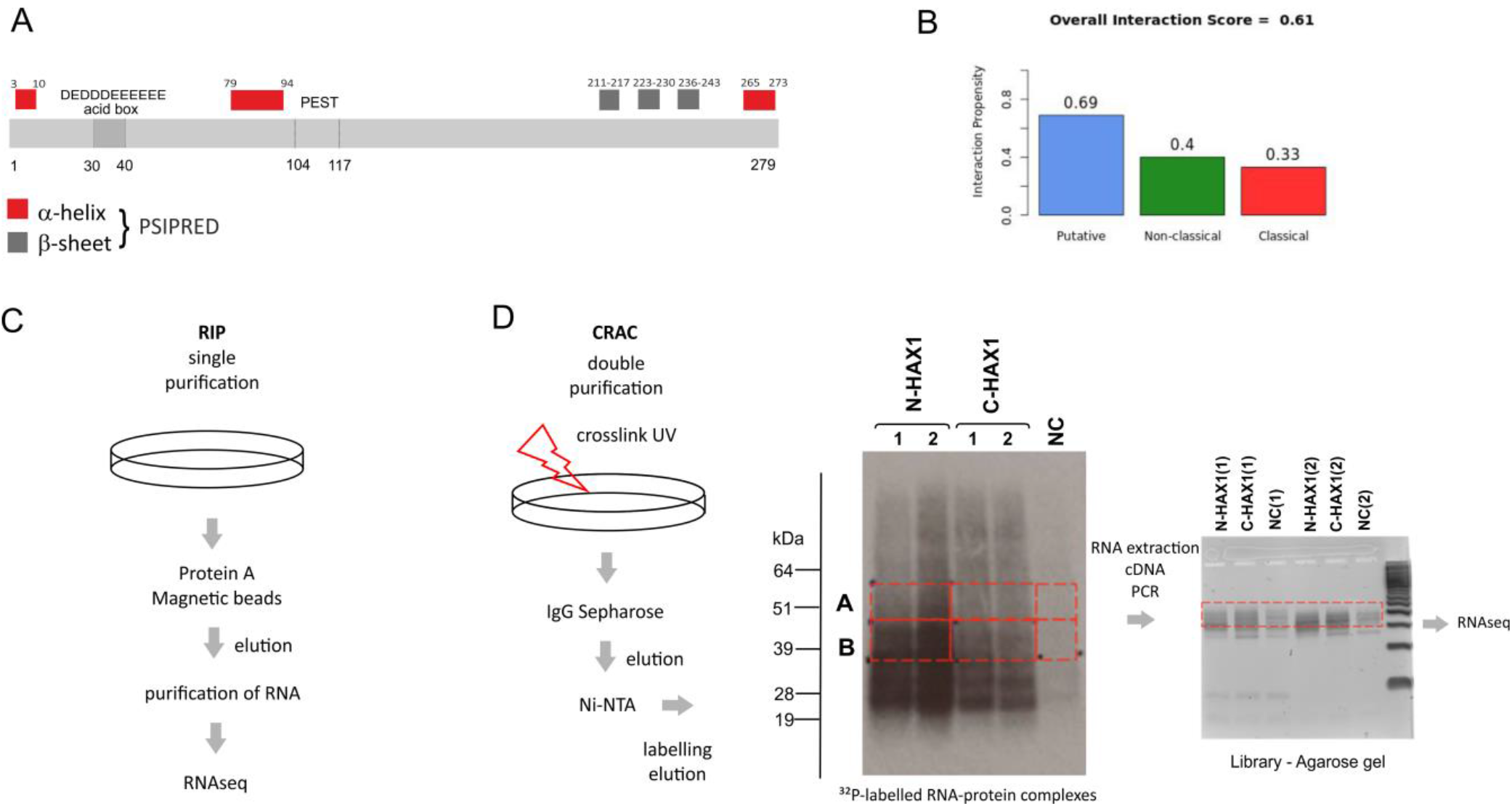
Experimental design of HAX1 –RNA interactome studies. A. HAX1 amino acid sequence with characteristic features and regions (acid box, PEST domain, structural elements predicted using PSIPRED [PSI-blast based secondary structure PREDiction]). B. CatRapid prediction of HAX1 RNA-binding propensity classifies it as a putative RBP, but the binding does not occur via classical elements C. Schematic of RIP experiments: no crosslink, single purification step with HAX1-specific antibody D. Schematic of CRAC experiments: UV crosslink of RNA targets, double purification.

#### cDNA library

Isolated RNA was incubated with SSIV reverse transcriptase (ThermoFisher) and a primer binding to 3’linker. cDNA was amplified using LA Takara Taq polymerase. PCR products were resolved on 3% Metaphor agarose gel (Lonza), and DNA fragments of sizes approximately 150-200 nt were isolated from the gel using Qiagen’s Gel Extraction Kit (Figure 1D). cDNA library was sequenced on the Illumina MiSeq platform in Edinburgh Genomics (The University of Edinburgh).

### 7. NGS Data Analysis (CRAC)

#### NGS results were analysed using algorithms

flexbar (preprocessing), tophat (genome mapping), and bedtools (analysis of genomic annotations). To identify HAX1 binding motifs, we extracted the coordinates of all HAX1 binding sites in mRNAs, and created a control dataset by randomly placing the coordinates of these binding sites on the same mRNAs using shuffleBed. We then used the STREME sequence motif discovery algorithm (minimum motif length 4 nt, maximum length 8 nt) to identify enriched motifs (https://meme-suite.org/meme/doc/streme.html, (Bailey 2021)). Genomic position of HAX1 binding identification was done using UCSC Genome Browser on Human Feb. 2009 (GRCh37/hg19) Assembly.

### 8. Gene Ontology and correlation analysis

Enrichment plots for RNA-seq data were generated by GSEA (Gene set enrichment analysis, (Mootha et al. 2003; Subramanian et al. 2005)) software. Table of transcripts, identified during RNAseq experiment, with associated number of counts per transcript (data obtained with HTSeq-count) was used as input data. Number of permutations was set to 1000 and permutation type was set to “gene_set”. Gene set database (version 7.2) used for analysis is included in respective enrichment plots titles. The other Gene Ontology enrichment analyses were performed using packages: Gene Ontology resource (http://geneontology.org) ; ((Ashburner et al. 2000); Gene Ontology 2021), Enrichr Ontologies (https://maayanlab.cloud/Enrichr/) (Kuleshov et al. 2016; Xie et al. 2021) and String11 Functional Enrichment Analysis for proteins with values/ranks (https://string-db.org) (Szklarczyk et al. 2021). Transcription factors were analyzed using Enrichr ENCODE and ChEA Consensus TFs from ChIP-X.

Correlation analysis with high-throughput data accumulated in TCGA (The Cancer Genome Atlas) database was performed using cBioPortal for Cancer Genomics, a platform for exploring multidimensional cancer genomic data (https://www.cbioportal.org/, (Cerami et al. 2012; Gao et al. 2013).

### 9. Transcription *in vitro*

Transcription *in vitro* was performed for 140 nt RPL19 fragment of CDS from exon 3 (primers: FW 5’-GGTGCATTATGCTTTCCCAGGTCAG-3’, REV 5’-CTATGCCCATGTGCCTGCCCTTC-3’) cloned into pGEM-T Easy vector in sense and antisense orientation. M13 fwd and rev primers were used in the PCR reaction for template generation. MEGAscript T7 transcription Kit (ThermoFisher Scientific) was used for *in vitro* transcription with T7 RNA polymerase, according to the manufacturers’ protocol. Transcripts were purified using MEGAclear Kit (ThermoFisher Scientific).

### 10. Microscale thermophoresis (MST)

MST experiments were performed using Monolith NT.115 (NanoTemper Technologies GmbH, Germany). Purified HAX1 protein (Proteintech, Ag27244, fused with His-tag) was labeled with RED-NHS 2nd Generation dye according to the supplied labeling protocol Monolith NT™ Protein Labeling Kit. A series of dilutions of ligand RNA (sense and antisense transcript) were prepared using buffer solution containing PBS with 0.2% Tween-20. The solution of labeled protein was mixed 1:1 with different concentrations of RNA strand yielding a final concentration of 50 nM of the protein and the ligand in a range of final concentrations between 10.8 µM and 0.000328 µM. After 5 min of incubation, the NT.115 premium capillaries (NanoTemper Technologies) were filled with the RNA/protein solution and thermophoresis was measured at a LED power of 100% and an MST power of 60% at RT. Each operation was controlled using MO.Control software. The K_d_ was determined by nonlinear fitting of the thermophoresis responses and EC50 was determined by Hill fitting model using the MO.Affinity Analysis v2.3 for both types of the calculations.

### 11. Western blot

Protein extracts were heat-denaturated (95°C) in Laemmli buffer (50 mM Tris/HCl, 0.01% Bromophenol Blue, 1.75% 2-mercaptoethanol, 11% glycerol, 2% SDS) and separated by 10–12% SDS/PAGE electrophoresis. Proteins were transferred to Immobilon-P PVDF membrane (Merck Millipore, MA, U.S.A). The membranes were incubated for 1h using 5% low-fat milk solution in 1X TBS (50 mM Tris-Cl, pH 7.5, 150 mM NaCl) as a blocking buffer and then overnight at 4°C in the same blocking solution containing one of the following antibodies: anti-HAX1 (rabbit, Proteintech 11266-1-AP), anti-RPL26 (rabbit, 1:5000, Abcam, ab59567). After washing (3×10 minutes in TBS), membranes were incubated for 2 h in room temperature with the adequate HRP-conjugated secondary antibody: goat anti-rabbit IgG (1:5000, Abcam, GB; cat. 97051) or goat anti-mouse IgG (1:10000, Abcam, GB; cat. ab97023). Membranes were developed using HRP detection kit WesternBright Quantum (Advansta, CA, U.S.A.; cat. K-12042).

### 12. qPCR

Quantitative PCR was performed as described (Grzybowska et al. 2013). Briefly, stable cell lines with *HAX1* WT and *HAX1* KO were subjected to Actinomycin D treatment (10 µg/ml). Cells were harvested in a designated time points and used for total RNA preparation (PureLink RNA mini kit; Invitrogen), followed by the treatment with recombinant DNase I (Roche). 1 µg of the obtained RNA was used for cDNA synthesis using Superscript III (Invitrogen). cDNA was quantified by quantitative PCR on an ABI Prism 7500 real-time PCR system using Power SYBR Green PCR Master Mix (Applied Biosystems, Life Technologies, Carlsbad, CA, USA) and primers amplifying a fragment of DHX37 transcript (forward 5’-CCCGATATCGAGAAAGCCTGG-3’; reverse 5’-CGTCCAGCACGTGAGATGAA-3’) and, as a reference, ACTB transcript (forward 5’-AGCCTCGCCTTTGCCGA-3’; reverse 5’-GCGCGGCGATATCATCATC-3’). The ΔΔCT method was used for calculating mRNA expression levels.

### 13. Sucrose gradient centrifugation

HL-60 cell lines (WT and *HAX1* KO#1 and #2) were grown and sub-cultured until achieving 6 T-75 or 3 T-175 flasks with cells at a density of 1×10^6^/ml. Cells were treated with cycloheximide (100 μg/ml) at 37°C for 10 minutes, harvested by centrifugation for 5 minutes at 500xg, 4°C, and washed 3 times with ice-cold PBS supplemented with 100 μg/mL cycloheximide. After final wash and complete removal of PBS, the cell pellet was resuspended in 0.75 ml of lysis buffer A [10 mM Tris-HCl pH 7.4, 12.5 mM MgCl_2_, 100 mM KCl, 0.5% Triton X-100 reduced, 2 mM DTT, 100 µg/mL CHX, 200 units SUPERaseIn™ RNase Inhibitor (20 U/μL; ThermoScientific), and cOmplete EDTA-free protease inhibitor (Roche)]. Cells were lysed by thorough pipetting and incubation for 15 minutes at 4°C on a rotating wheel. Lysates were aspirated into 1 ml syringe, passed through a 26G needle seven times and then centrifuged for 10 minutes at 16000xg, 4°C. RNA concentration in clarified cytoplasmic extracts thus obtained was measured using Nanodrop 2000c (ThermoScientific). 14-20 OD_260_ units of cytoplasmic extract in 500 μl of lysis buffer was layered on top of 10-50% linear sucrose gradients, prepared using ÅKTA Purifier FPLC system and 0.22 µm-filtered sucrose solutions in polysome buffer [20 mM Hepes-KOH pH 7.4, 12.5 mM MgCl_2_, 100 mM KCl; 2 mM DTT, 100 µg/ml CHX, and cOmplete EDTA-free protease inhibitor], and ultracentrifuged for 3h15’ at 36000 rpm, 4°C in SW-41Ti rotor (Beckman Coulter). Subsequently, 0.5 ml fractions were collected from gradient by pumping 60% sucrose solution in polysome buffer to the bottom of tubes and OD_254_ was monitored on ÅKTA Purifier.

## RESULTS

### 1. Different high-throughput analyses of HAX1 binding targets consistently reveal its involvement in ribosome biogenesis and translation

HAX1 is predicted to contain few secondary structure elements, while most of its sequence is predicted to be disordered and there is no conventional RNA-binding domain (Figure 1 A, (Balcerak et al. 2017; Larsen et al. 2020)). Nevertheless, RNA-binding propensity of HAX1 was reported previously (Al-Maghrebi et al. 2002; Sarnowska et al. 2007). Moreover, three different RNA-binding prediction algorithms suggest that HAX1 may bind RNA: RNAPred (the score for putative RBD: 0.58 SVM [support vector machine], cutoff: 0.5, https://webs.iiitd.edu.in/raghava/rnapred/), RBPPred (the score for putative RBD: 0.928 SVM, cutoff: 0.5, http://rnabinding.com/RBPPred.html) (Zhang and Liu 2017) and CatRapid (the score for putative RBD: 0.61, threshold 0.5, http://s.tartaglialab.com/page/catrapid_group) (Livi et al. 2016) (Figure 1B). Algorithms comparing three-dimensional structure cannot be used, since HAX1 has no homologs, is mostly disordered and has not been crystalized.

To verify and globally characterize RNA-binding propensity of HAX1, we employed two independent, high throughput screens (RIP-seq and CRAC) for HAX1 RNA targets (Figure 1C,D). RIP-seq is relatively straightforward, with one purification step and without the crosslinking step and thus it yields more background noise (>5000 hits). Conversely, CRAC involved RNA crosslinking and two purification steps, thus, the results were more specific and enabled the analysis of the bound RNA regions. Both approaches unanimously revealed enrichment in RNAs linked to ribosome biogenesis, translation, co-translational targeting to the membrane and RNA processing.

#### 1.1. RIP-seq results

RNA target identification was performed in HL-60 human leukemia cell line, because promyelocytes represent the most affected cell type in patients with recessive mutation in both HAX1 alleles (severe neutropenia), so analyzing the role of HAX1 in these cells seems to be the most physiologically relevant. HAX1-associated transcriptome was identified by employing the next generation sequencing after RNA immunoprecipitation (RIP-seq) of HAX1-complexes. Transcripts were classified as positive targets using an adjusted p-value cutoff of 0.05. Enrichment analysis (Enrichr) of the obtained dataset indicates involvement in ribosome biogenesis and RNA processing, including a strong category of targets involved in rRNA processing (Biological Process). Molecular Function analysis points mostly to RNA binding, while Cellular Component indicates nucleolus and preribosome (Figures 2A and S2A, supplementary data file S1). Plotting enrichment against false discovery rate (FDR) for GO terms obtained by Gene Ontology Enrichment Analysis (Panther) in Biological Process category revealed that terms related to ribosome (including ribosome biogenesis and rRNA processing) and translation have high significance (high enrichment, low FDR, Figure 2B). Most of the RNA targets obtained in RIP analysis represent mRNAs (Figure 2C). Figure 2D represents the RIP-seq coverage (X-axis; FPKM, fragments per kilobase per million, to compensate for the length of the transcript) plotted against log2 enrichment in RIP-seq (RIP-seq_HAX1/RIP-seq_IgG). Red points above (or below) the horizontal axis represent transcripts that are significantly enriched (or depleted) in the HAX1 pulldown, compared to their total abundance in cells.

**Figure 2.**
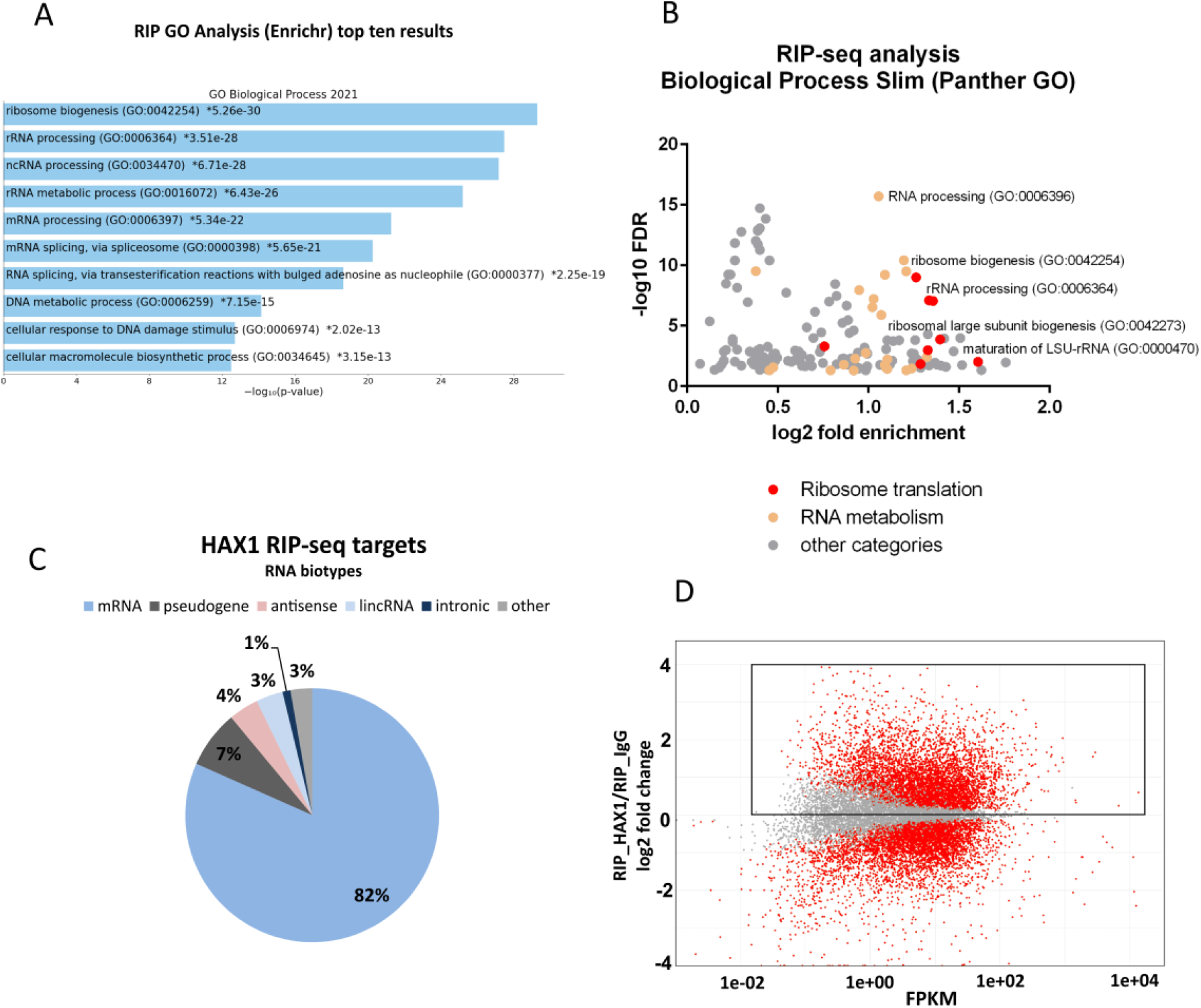
Identification of HAX1-associated RNA targets by RIP-seq analysis in HL-60 cells. A. Functional annotation enrichment of HAX1 target genes. Enriched Gene Ontology terms (Enrichr Ontologies: Biological Process) presented in bar plot, the length of each bar corresponds to statistical significance of the enrichment (-log10 from adjusted p-value). Top ten terms are presented. B. Enriched Biological Process terms (Panther GO) plotted with fold enrichment (log2) against False Discovery Rate (-log10). The most reliable results are in the right upper corner. Categories associated with ribosome biogenesis, rRNA processing and translation are marked red. Categories associated with RNA metabolism (other than rRNA) are marked light orange. Detailed description of GO terms in Supplementary filel S1. C. Distribution of RNA classes among HAX1 RIP targets. D. MA-plot showing RIP targets counts (X-axis) calculated as fragments per kilobase per million (FPKM); Y-axis: log2FC(RIP_HAX1/RIP_IgG), red-significant, grey-not significant. Internal frame indicates positive log2 FC (physiologically relevant as possible binding targets).

#### 1.2. CRAC results

CRAC analysis was performed in HEK293FlpInTRex human embryonic kidney cell line modified to overexpress *HAX1* upon doxycycline induction. This cell line was used because it has been standardized for CRAC experiments. Two different cell lines were generated with a tag added on 5’ or 3’ end of HAX1 protein coding sequence and compared to negative control cell line. Results of these complementary experiments were pooled in an overlapping list of 223 targets (log2FC[CRAC_HAX1/CRAC_NC]>0), which were analyzed further (supplementary data file S2). Enrichment analysis (Enrichr) revealed involvement in translation and ribosome assembly, including co-translational targeting to membrane, rRNA processing and ribosome biogenesis (Biological Process). Molecular Function analysis points mostly to RNA binding, while Cellular Component indicates various categories linked to ribosome (Figures 3A and S2B, supplementary data file S2). Enrichment values plotted against false discovery rate (FDR) for GO terms obtained by Gene Ontology Enrichment Analysis (Panther) revealed that similar terms as in RIP (involved in translation, rRNA processing and ribosome biogenesis) are of the highest significance (Figure 3B). Analyses performed separately for with HAX1 tagged on C or N terminus (respectively 1637 and 767 targets) revealed enrichments with the same terms as pooled analysis (Figure S2C,D, supplementary data file S2).

**Figure 3.**
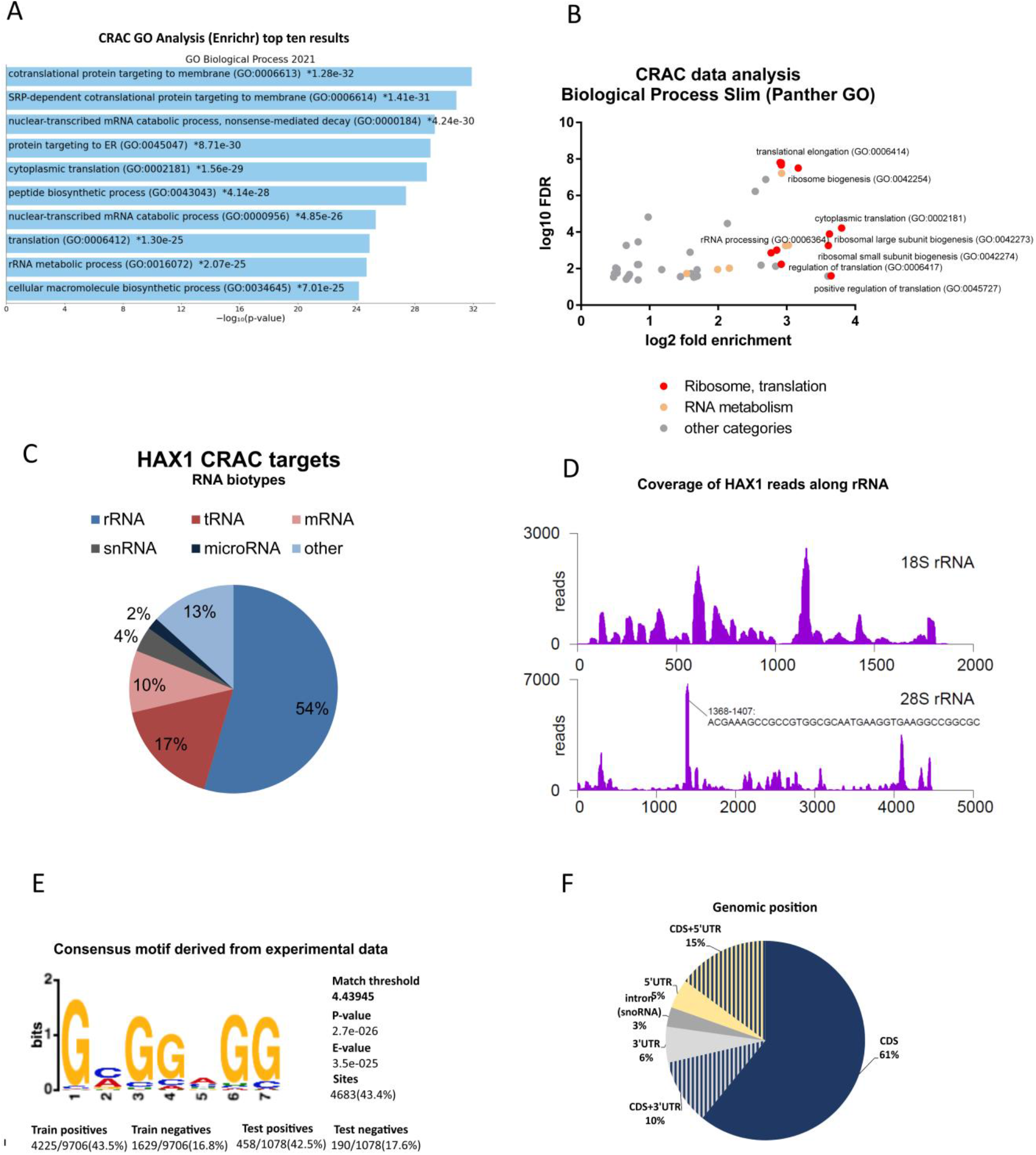
Identification of HAX1-associated RNA targets by CRAC analysis (HEK293 cells). A. Functional annotation enrichment of HAX1 target genes (overlapping target pool from analyses with N-terminal and C-terminal tags). Enriched Gene Ontology terms (Enrichr Ontologies: Biological Process) presented in bar plot, the length of each bar corresponds to statistical significance of the enrichment (-log10 from adjusted p-value). Top ten terms are presented for each category. B. Enriched Biological Process terms (Panther GO) plotted with fold enrichment (log2) against False Discovery Rate (-log10). The most reliable results are in the right upper corner. Terms associated with ribosome biogenesis, rRNA processing and translation are marked red. Terms associated with RNA metabolism (other than rRNA) are marked light orange. Detailed description of GO terms in supplementary data file S2. C. Distribution of the RNA species in HAX1 targets (CRAC) D. Coverage of HAX1 CRAC reads along small (18S) and large (28S) ribosomal subunits. E. Consensus motif identified by STREME in the HAX1-targets dataset (combined experimental data from C- and N-tagged HAX1). The E-value is the p-value multiplied by the number of motifs reported by STREME. F. Genomic position of HAX1 binding in CRAC-identified targets (188 targets overlapping in C- and N-tagged datasets) established using UCSC Genome Browser on Human Feb. 2009 (GRCh37/hg19) Assembly

Transcript biotypes analysis indicates high proportion of rRNA and tRNA (respectively 54% and 17%) with mRNA representing 10% of CRAC targets (Figure 3C). Detailed analysis of the coverage of HAX1 reads along small (18S) and large (28S) ribosomal subunits shows high reads for the region of 1368-1407 in the large subunit, indicating possible interaction site (Figures 3D and S3).

CRAC method enables the assessment of the enrichment in recurring sequence patterns (motifs) in an analyzed experimental dataset. Motif finding was performed using STREME (The MEME Suite; Motif-based sequence analysis tools) resulting in a characterization of a guanine-rich motif for the pooled C and N-terminally tagged datasets (Figures 3E and S4).

The analysis of genomic position of detected mRNA targets (188 targets from the pooled C and N datasets) revealed prevalent HAX1 binding to the coding sequence (CDS) region and substantial number of targets located in both CDS and 5’ or 3’ UTRs. Only 14% of targets mapped in non-coding regions, 3% of which represented introns. Interestingly, while introns alone represent about 3% of all target positions, introns plus CDS represent about 11% and 47% of these intronic targets encode snoRNA (supplementary data file S2).

#### 1.3. Comparison of RIP-seq and CRAC results

In both approaches enriched GO terms encompassed ribosome biogenesis, translation, rRNA processing and RNA processing in general. Enrichment in the same categories was detected for the group of 72 targets, overlapping in both screens, these categories were also the most probable (Figure 4A,B). For this overlapping group, categories associated with translation and ribosome biogenesis were shown to form a tight cluster in String 11 analysis (Figure 4C), with a high proportion of ribosomal proteins and ribosome assembly factors. Analysis of the group of the overlapping targets in RIP and CRAC revealed that while for the total numbers the overlap was tiny (1.2%), when specific group of targets were compared (characterized by Enrichr, supplementary data file S3) the overlap increased to 12.4% and 14.4 % respectively for protein coding transcripts involved in ribosome biogenesis and translation (Figure 4D). These results point to the probable role of HAX1 in ribosome assembly and translation.

**Figure 4.**
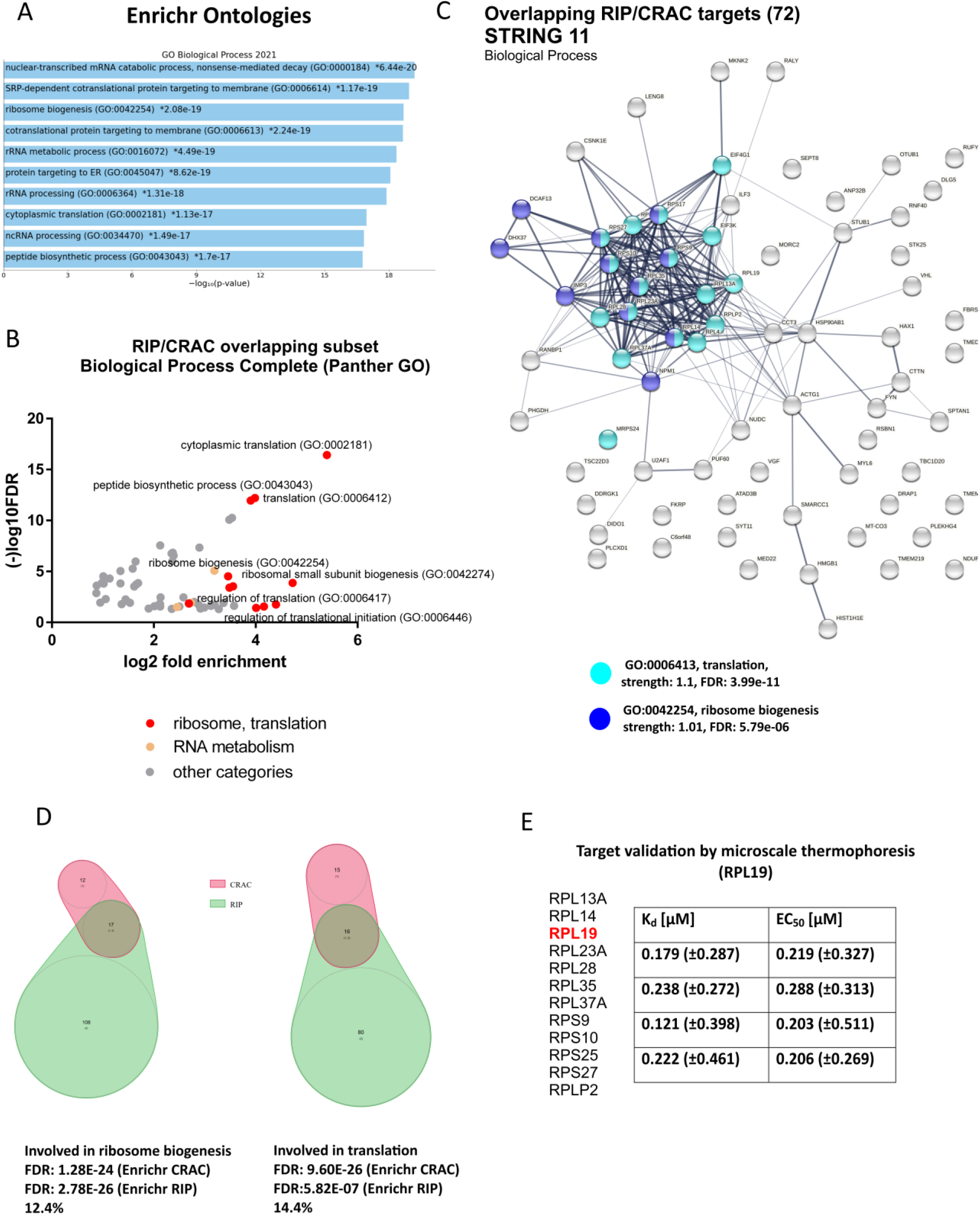
Characterization of the overlapping CRAC-RIP subset of HAX1 mRNA targets. A. Functional annotation enrichment of the RIP-CRAC overlap of HAX1 target genes (Enrichr Ontologies, top ten terms) maintains high enrichment in terms associated with ribosome biogenesis, translation, rRNA processing (Biological Process) B. Enriched Biological Process terms found for the overlapping subset (Panther GO) plotted with fold enrichment (log2) against False Discovery Rate (-log10). C. String11 analysis of the overlapping CRAC/RIP targets reveals a cluster of proteins involved in translation and ribosome biogenesis D. Overlapping targets in RIP and CRAC results for subsets of transcripts categorized by Enrichr in GO terms involved in ribosome biogenesis (12.4% overlap) and translation (14.4% overlap). The overlap is significantly higher in these subsets than in total results without categorization (1.2 % overlap between RIP and CRAC). E. Target validation by microscale thermophoresis (MST) performed for RPL19 *in vitro* transcript (sense). From 12 transcripts encoding ribosomal proteins present in the overlapping RIP/CRAC target list (left), a fragment of RPL19 coding sequence was selected for validation. Interaction with purified, fluorescently labeled HAX1 protein was confirmed in 4 independent measurements by thermophoresis in different conditions (MST power medium to high). For each measurement dissociation constant (K_d_) and half maximal effective concentration (EC_50_) was assigned. Interaction with antisense transcript (negative control) was not observed.

#### 1.4. Validation of selected target by microscale thermophoresis

Microscale thermophoresis (MST) was performed for the *in vitro* transcript containing a fragment of the coding sequence of RPL19 ribosomal protein. This target was selected due to its relatively high rank in the results obtained by both methods (RIP and CRAC). Specific sequence was based upon CRAC identification of the binding region within exon3. Titration of the target RNA to a constant concentration of the fluorescently labeled HAX1 protein allowed to determine ligand dissociation constants (K_d_), and systematically compare their binding affinity. Half maximal effective concentration (EC50) was calculated by the Hill fit model. The results reveal binding for the sense transcript, but not for the antisense negative control (Figure 4E).

### 2. *HAX1* KO affects expression profile of HL-60 cells

To assess the impact of HAX1 on HL-60 transcriptome, two independent cell lines with *HAX1* CRISPR/Cas9 knockout were generated (*HAX1* KO#1, KO#2, Figure EV1) and used in RNA-seq experiment to compare expression profiles with *HAX1* WT cells. Statistical significance was assigned only for genes differentially expressed in both *HAX1* KO cell lines vs. WT and in the same direction (p-value cutoff 0.05). Gene ontology analysis revealed that HAX1 knockout affects the expression profile of HL-60 cells in several terms of Biological Process (Figure 5A-C). Weighted analysis of RNA-seq results (String 11, functional enrichment analysis for proteins ranked according to fold change [RNA-seq_*HAX1* KO/RNA-seq_WT]) allows to distinguish two main groups: GO terms linked to ribosome biogenesis, rRNA processing and translation (including mitochondrial translation) and terms linked to energy generation in mitochondria: respiratory electron transport chain and oxidative phosphorylation (described in detail in supplementary data file S4). *HAX1* knockout affects significantly the expression of 2344 genes (1158 upregulated and 1186 downregulated in KO). As indicated in Figure 5C, genes linked to ribosome biogenesis, translation and energy generation in mitochondria tend to be downregulated in *HAX1* KO cells.

**Figure 5.**
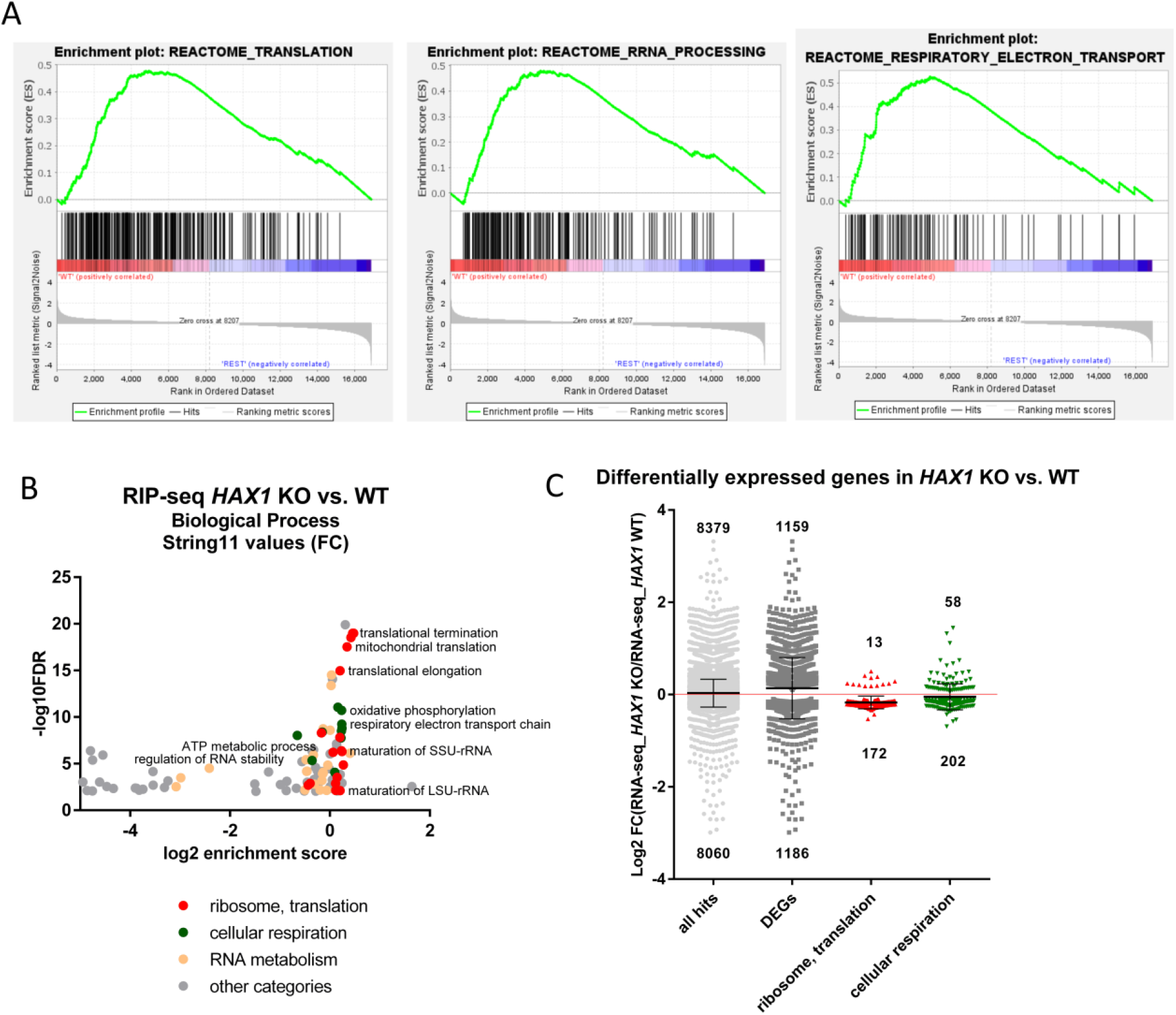
*HAX1* KO affects expression profile in HL-60 cells. A. Enrichment plots generated by GSEA analysis for the chosen categories (translation, rRNA processing, respiratory electron transport) B. Enrichment in GO Biological Process terms for genes differentially expressed in *HAX1* KO vs. HAX1 WT (String11 weighted analysis with log2[RNA-Seq_*HAX1*_KO/RNA-Seq WT] as values) indicates involvement in translation (including mitochondrial translation), rRNA processing/ribosome biogenesis and energy generation in mitochondria. Detailed description of GO terms in supplementary data file S4. C. Scatter plot showing differentially expressed genes in *HAX1* KO. From the total 1186 transcripts significantly downregulated in KO, 172 are associated with translation and 202 with energy generation, while from 1158 transcripts significantly upregulated in KO 13 and 58 respectively are associated with these categories. p-value cutoff: 0.05. 14 outliers were omitted. Statistical significance of downregulation of the transcripts involved in ribosome biogenesis and translation and cellular respiration was assessed by Chi-Square test and was proved to be high (p-value=5.5E-20 and 1.2E-05 respectively). For the other two groups (all mapped results and all DEGs) there was no statistical significance associated with downregulation.

### 3. Correlation analysis of *HAX1* expression in cancer databases reveals differences in the same GO terms as transcription profiling in HL-60 cells

Correlation of *HAX1* expression with other genes can be analyzed in expression databases created using high-throughput methods applied to patients’ samples from many different neoplasms. Neoplasms selected for correlation analysis were chosen as corresponding to HL-60 cell line (leukemia) and another hematologic neoplasm (lymphoma) or common types of cancer of epithelial origin (breast, cervical cancer).

Co-expression analysis was performed using cBioPortal for Cancer Genomics with data from TCGA (The Cancer Genome Atlas, PanCancer Atlas) for the four neoplasms (Cervical Cancer - 297 patients, Breast Cancer – 1084 patients, Acute Myeloid Leukemia – 200 patients and Diffuse Large B-cell Lymphoma – 48 patients). Gene lists with the Spearman’s rank correlation coefficients were used in String 11 gene ontology analysis (with Spearman’s coefficients as values/ranks) and obtained Biological Process terms were plotted in Figure 6A-D, showing that terms with the highest enrichment and the lowest false discovery rate are similar as in expression profiling obtained in HL-60 cells (RNA-seq). The most enriched and probable terms revealed by weighted analysis of genes which expression significantly correlates with *HAX1* expression include those involved in translation (co-translational protein targeting to membrane, mitochondrial translation, translational elongation, translational initiation, cytoplasmic translation), ribosome assembly and rRNA processing, but also energy generation in mitochondria (respiratory electron transport chain, oxidative phosphorylation, cellular respiration) and RNA processing in general. These enrichments were observed in all analyzed neoplasms. For details see supplementary data file S5.

**Figure 6.**
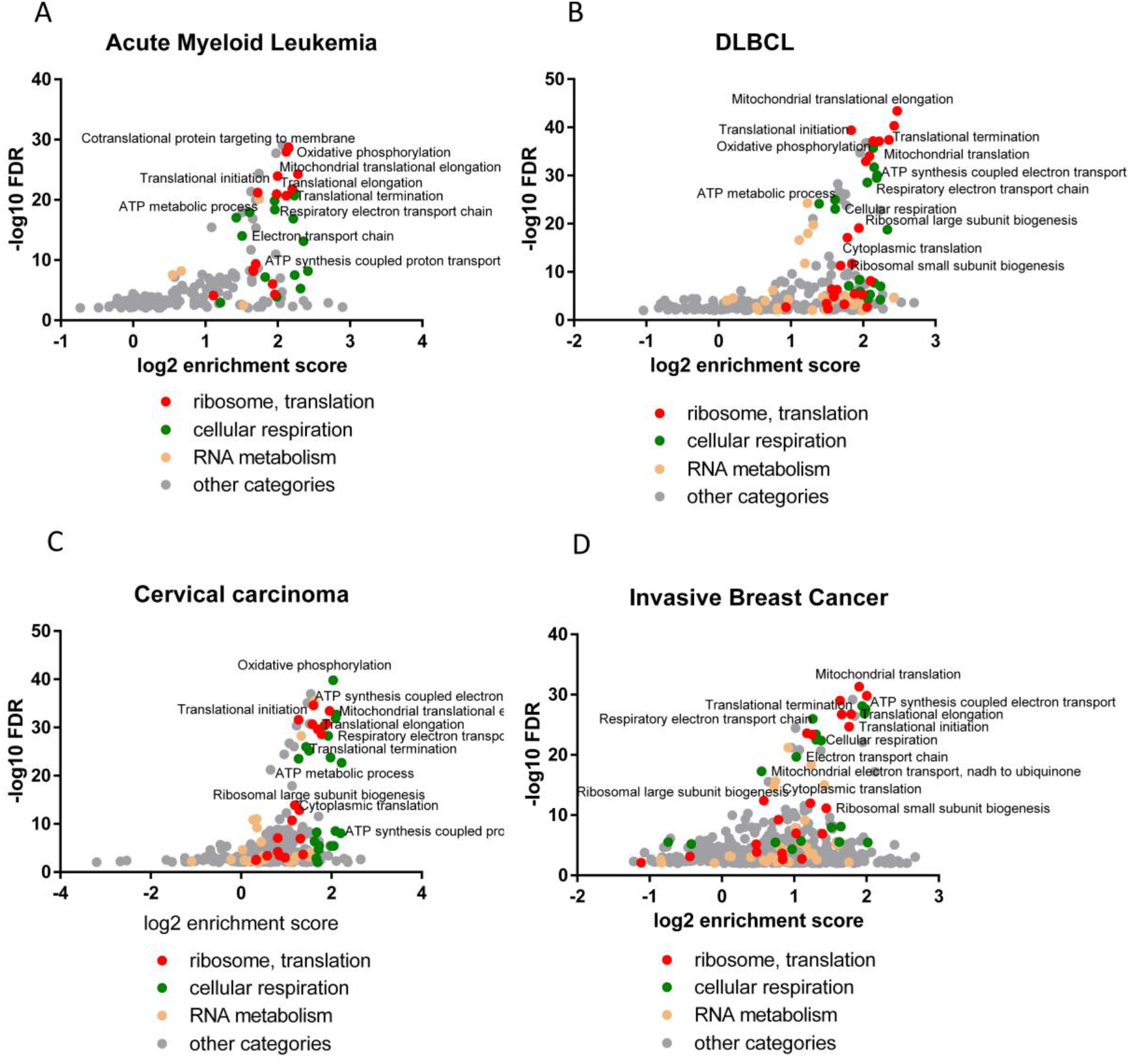
Subset of genes whose expression correlates with HAX1 in the four different neoplasms display enrichment in the same biological processes as shown for *HAX1* KO. Correlation analysis was performed using cBioPortal Cancer Genomics. Clinical data obtained from TCGA database. Enrichment analysis was performed using String11 with correlation coefficients as values. A. Acute Myeloid Leukemia (AML, 200 patients) B. Diffuse Large B-cell Lymphoma (DLBCL, 48 patients) C. Cervical cancer (297 patients) D. Invasive breast cancer (1084 patients) C. Detailed description of genes, Spearman coefficients and GO terms in supplementary data file S5.

Similar enrichments were also observed in other types of cancer (renal cell carcinoma, glioblastoma,data not shown).

### 4. RIP-seq and RNA-seq comparison suggests that HAX1-binding may stabilize a subset of transcripts involved in ribosome biogenesis

RIP-seq analysis and RNA-seq for *HAX1* KO/WT were performed in the same HL-60 cell line, thus the comparison should reveal if RNA binding has an effect on RNA stability.

Figure 7A represents the same MA plot as on Figure 2D, showing RIP-seq results with superimposed information about transcripts differentially expressed in *HAX1* KO cells (RNA-seq_*HAX1* KO/RNA-seq_WT): marked in blue (downregulated) and in red (upregulated). DEGs representation was trimmed to the relevant RIP results (internal frame). This combined analysis of the results of RIP-seq and RNA-seq reveal that transcripts differentially expressed in RNA-seq are also enriched in HAX1 RIP-seq, suggesting that HAX1 binding affects the expression. Interestingly, transcripts downregulated in *HAX1* KO concentrate in the most significant part of the plot, with the highest counts, suggesting that HAX1 binding may cause stabilization.

**Figure 7.**
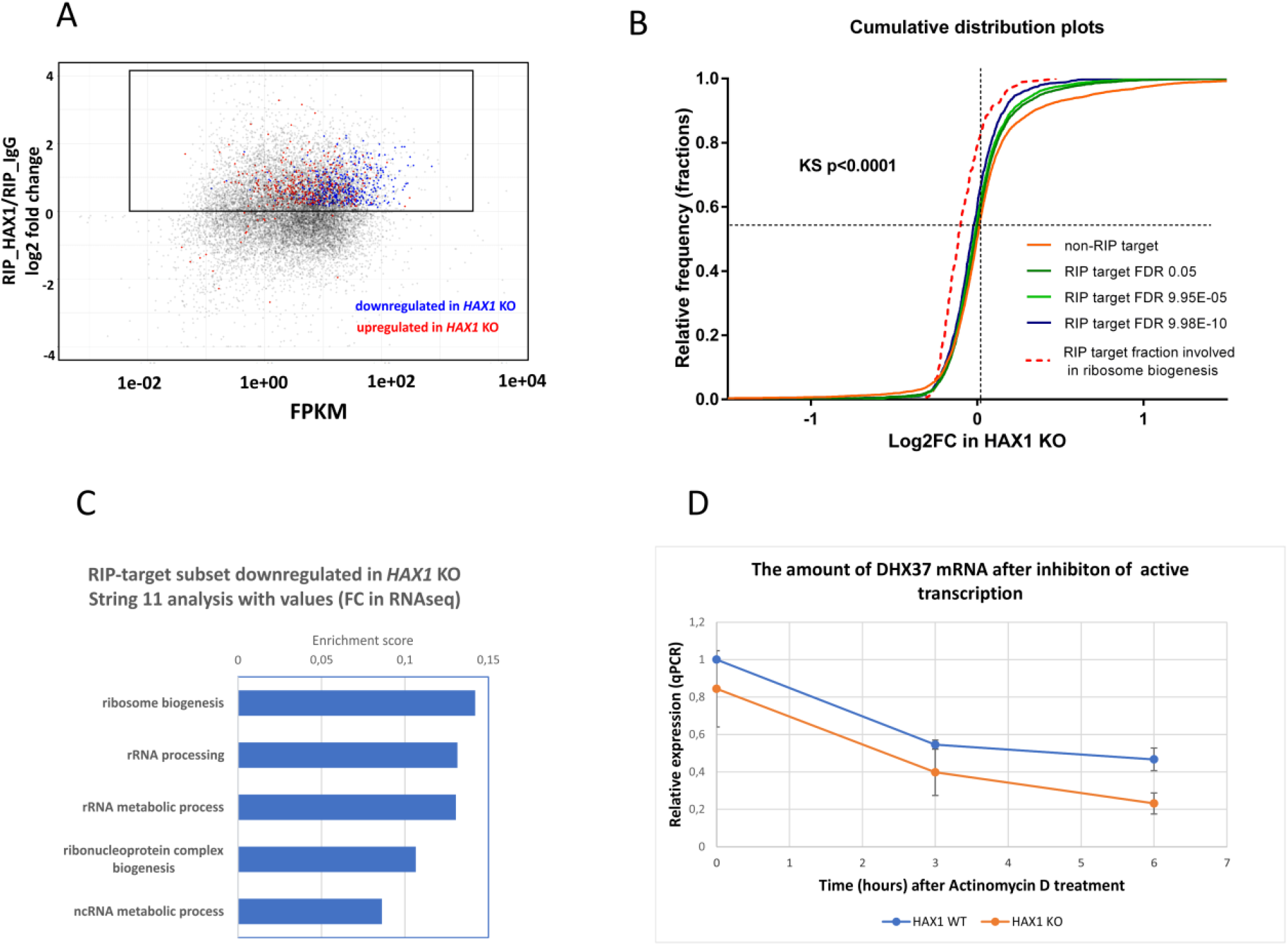
HAX1 binding to RNA targets may be responsible for changes in the expression profile A. MA-plot showing RIP targets counts, same as in Figure 2D, with superimposed RNA-seq results, marking DEGs; X-axis: fragments per kilobase per million (FPKM); Y-axis: log2FC(RIP_HAX1/RIP_IgG), RIP targets with significantly downregulated expression upon *HAX1* KO (RNA-seq_*HAX1* KO/RNA-seq_WT) marked blue, RIP targets with significantly upregulated expression marked red, remaining targets marked grey (shadowed for clarity). Internal frame indicates relevant RIP results with positive log2 FC, RNA-seq results trimmed to internal frame. B. Comparison of distribution of FC changes caused by *HAX1* knockout (data obtained in RNA-seq) in the subset of transcripts present among targets identified by RIP (RIP target) and not identified by RIP (non-RIP target). Three different RIP-target subsets were compared with cutoffs determined according to the decreasing FDR, showing a shift towards lower FC values (RIP_HAX1/RIP_IgG) in the subsets with lower FDR. Dotted line: a subset of transcripts involved in ribosome biogenesis, identified by String11 weighted analysis for RIP-target transcripts (RIP_HAX1/RIP_IgG FC as values). Significance calculated by Kolmogorov-Smirnov test C. Enrichment in GO terms assessed by String11 weighted analysis with FC values from RNA-seq for a subset of downregulated transcripts. Analogous analysis for upregulated subset produced no results. D. DHX37 mRNA degradation is more dynamic in *HAX1* KO cells. Cells were treated with Actinomycin D (10 µg/ml). Relative expression was quantified by qPCR in designated time points. The experiment was performed in 4 biological repeats (n=4).

To further analyze this effect, RNA-seq fold change (RNA-seq_*HAX1*_KO/RNA-seq_WT) distributions of transcripts representing HAX1 targets (RIP target subset) and the rest of transcripts (non-RIP target subset) were compared (Figure 7B), showing that changes in distribution towards downregulation of RIP-associated transcripts in *HAX1* KO increase with the increasing probability of the HAX1-specific RNA interaction (lower FDR) suggesting that RIP targets are less stable. The effect, while statistically significant is not ubiquitous and does not affect all RIP targets equally, suggesting some other layers of regulation. Distribution analysis performed only for transcripts involved in ribosome biogenesis (dotted line) shows much more substantial shift towards lower FC values (log2[RNA-seq_*HAX1*_KO/RNA-seq_WT]). Additionally, the fraction of transcripts involved in ribosome biogenesis increases when RIP results are analyzed with the same FDR cutoffs in weighted analysis (String11, Biological Process), with FC from RNA-seq as values (Figure S5A).

Interestingly, HAX1-binding seem to be limited to the transcripts linked to ribosome biogenesis and not include transcripts linked to oxidative phosphorylation/respiratory electron chain, which were also significantly downregulated in *HAX1* KO, indicating a different mode of regulation for this group of transcripts. Additionally, mRNA RIP targets (RIP target subset) along with their respective FCs in RNA-seq were analyzed in weighted analysis (String11, values, Biological Process), separately for downregulated and upregulated transcripts, revealing an enrichment in ribosome biogenesis and rRNA processing (GO:0042254 and GO:0006364) for the group of downregulated transcripts, while the same analysis of the upregulated group did not reveal any enrichment (Figure 7C).

Moreover, the possibility of direct stabilization of transcripts by HAX1-binding was tested for DHX37 mRNA (encoding DEAH-box helicase 37 involved in ribosome biogenesis and translation initiation). DHX37 mRNA was selected from transcripts downregulated in *HAX1* KO and simultaneously from RIP/CRAC analysis, as a potential HAX1 target, with a binding region within the coding sequence of the transcript. Transcription was inhibited by Actinomycin D treatment and subsequent mRNA degradation was analyzed by qPCR in *HAX1* WT and *HAX1* KO cell lines, indicating more dynamic degradation in HAX1-deficient cell line (Figure 7D).

To test the possibility that HAX1 effect on transcriptome may not be direct, but may instead consist in influencing transcription factors (TFs) and propagating the effect to the group of genes regulated by these factors, TFs potentially regulating the subset of genes for which the expression has changed in *HAX1* KO were identified using Enrichr package (Transcription, ENCODE and ChEA) and juxtaposed with the ranked list of RIP targets. TFs of the highest rank (FC, FDR) and the highest Combined Score provided by Enrichr are listed in Figure S5B (higher panel). YY1 and USF1 are the only TFs with FC >2, but MYC, which has the highest Combined score, while not highly ranked itself, should be considered because MYC-binding proteins are also among RIP targets, with MYCBP of FC>3. Genes regulated by all these TFs are mostly downregulated in *HAX1* KO (Figure S5B, lower panel). However, the expression of all these TFs is not changed at the mRNA level, undermining the possibility of the indirect regulation via TFs.

### 5. *HAX1* KO affects 40S:80S ratio

To assess physiological effect of HAX1 deficiency in the context of ribosome biogenesis, ribosome sedimentation in sucrose density gradient was performed for *HAX1* WT and *HAX1* KO HL-60 cells. Ribosomal subunit stoichiometry ratios 40S:60S and P/M (polysome-to-monosome) did not reveal significant changes between WT and KO cell lines, but the ratio of 40S to 80S revealed that the peak corresponding to the small subunit is increased in *HAX1* KO compared to WT (Figure 8A). This effect is significant for both *HAX1* KO cell lines, while the overall subunits:monosome ratio (40S+60S:80S) is significantly increased only for one KO cell line (Figure 8B).

**Figure 8.**
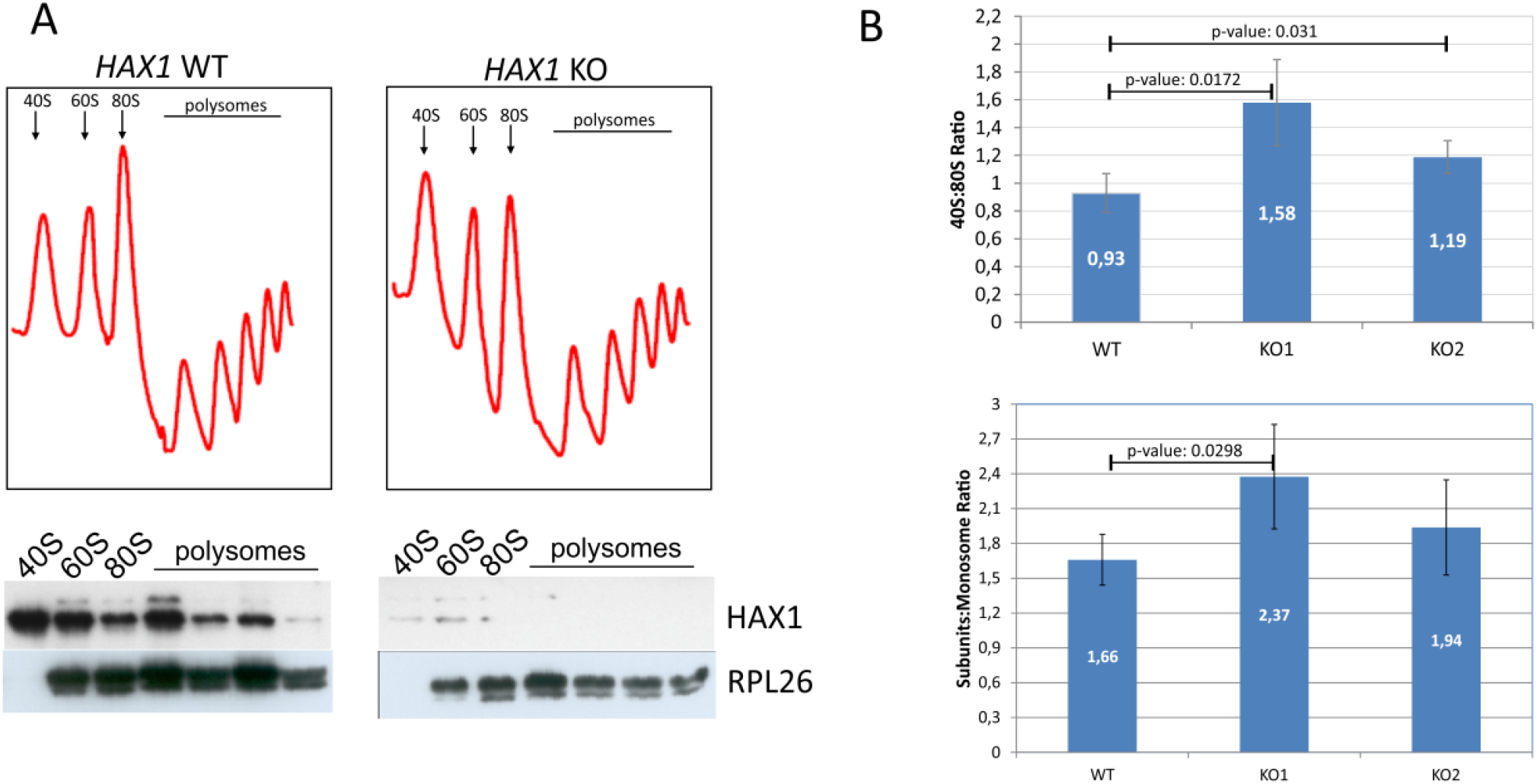
*HAX1* KO affects subunit:monosome ratio. A. Absorbance profiles at 254 nm of sucrose gradient (10–50%) sedimentation of HL-60 cell extracts. The three leftmost peaks include ribosomal subunits (40S and 60S) and non-translating monosomes (80S). The remaining peaks represent polysomes. B. Quantitative analysis of the sedimentation profiles indicates that free subunits are more abundant in *HAX1* KO in relation to monosome. Area under each peak was quantified using ImageJ (n=6 in each sample), statistical significance determined with unpaired t test.

## DISCUSSION

In this manuscript we report a comprehensive analysis of HAX1 RNA-interactome by two independent, high-throughput methods, which both suggest HAX1 involvement in the regulation of transcripts controlling ribosome biogenesis, rRNA processing and translation. The two methods differ in complexity and specificity (Figure 1A), and experiments were performed in two different cell lines (HL-60 promyelocytic cell line, selected due to the strong effect of HAX1 inactivation in these cells and HEK293 embryonic kidney cells with adrenal endocrine characteristics, selected for technical reasons). Nevertheless, both approaches yielded an enrichment in similar GO terms in Biological Process category (ribosome biogenesis, rRNA processing, translation). Molecular Function analysis indicated high proportion of transcripts implicated in RNA binding, for both RIP and CRAC. Cellular Component analysis indicated nucleolus and preribosome for RIP targets, and various ribosomal categories for CRAC targets (cytosolic ribosome, large and small ribosomal subunits). Targets common for both methods are involved in ribosome biogenesis and translation initiation, and 72 targets that constitute RIP/CRAC overlap are mostly composed of transcripts encoding ribosomal proteins and ribosome biogenesis/translation factors.

Distribution of RNA biotypes detected in both methods was different, with RIP targets predominantly consisting of mRNAs and CRAC targets with high proportion of rRNAs and tRNAs. These differences may stem from the fact that in CRAC the protein of interest was overexpressed and RNA targets were UV-crosslinked to it, which may shift the distribution towards less stable, transient interactions, comparing to endogenous RIP.

The next important question was if HAX1 RNA targets expression has changed upon *HAX1* knockout. This comparison was conducted for HL-60 cells, since this cell line was used in both, RIP and RNA-seq expression profiling for *HAX1* KO vs. WT. The results indicate partial overlap between HAX1 RNA targets and mRNAs downregulated in *HAX1* KO and the overlap pertained to transcripts involved in ribosome biogenesis and translation, and only to very small extent to transcripts involved in energy generation in mitochondria, also significantly downregulated in *HAX1* KO. This result suggests that only the subset involved in ribosome biogenesis and translation is regulated via direct HAX1 binding and subsequent mRNA stabilization. The other detected changes (especially the downregulation of a very important subset of transcripts involved in energy generation) must be therefore regulated by a different mechanism.

Further support for the hypothesis of HAX1 transcript-stabilizing role is provided by quantitative assessment of the degradation of DHX37 mRNA. This RNA helicase is involved in ribosome biogenesis (Choudhury et al. 2019) and its mRNA represents one of the top RIP/CRAC targets downregulated in *HAX1* KO cells. Quantification of the DHX37 transcript degradation revealed more dynamic degradation in *HAX1* KO cells, suggesting stabilization by HAX1..

Analysis of genomic position of RNA targets obtained using CRAC method revealed the prevalence of coding sequence (CDS) regions, which is not typical for the regulatory RNA sequence and not consistent with the genomic position of previously characterized HAX1 binding regions (3’UTRs). However, new high-throughput analyzes demonstrated that binding to the CDS is not as uncommon as previously thought and may have a role in the regulation of mRNA stability (Grzybowska and Wakula 2021). Interestingly, this stabilization should refer to the situation when mRNA is not actively translated, thus it is not covered and protected by ribosomes and susceptible to endonuclease attack, like in case of protein CRD-BP, which binds to c-myc mRNA, protecting it (Lemm and Ross 2002). Moreover, CDS binding was also observed for protein FMRP and linked to the recruitment of the APP mRNA to processing bodies (P-bodies), which was proposed to restrict translation (Lee et al. 2010). In line with this observation, we reported previously that HAX1 was observed to co-localize with P-body marker, Dcp1 (Zayat et al. 2015), pointing to its possible role in transcript stabilization during storage.

Interaction with one of the targets within CDS reported by both methods (RPL19) was confirmed by microscale thermophoresis, with the obtained dissociation constants indicating relatively weak binding. These values suggest transient, regulatory interaction, typical for intrinsically disordered proteins. Interestingly, K_d_ values previously reported for HAX1 binding to 3’UTR are lower (Sarnowska et al. 2007), indicating different strength of interaction and, possibly, different mode of binding for CDS and 3’UTR regions. Similar phenomenon was also observed for GLD-1 and FMRP proteins, involved in the regulation of translation (Grzybowska and Wakula 2021).

Analysis of the possibility of indirect regulation mediated by transcription factors indicated that such regulation is improbable, since the expression of TFs themselves is not changed in *HAX1* KO. This hypothesis cannot be however totally dismissed, since TFs-encoding transcripts may be differentially translated or their protein product may be degraded in *HAX1* KO cells, resulting in differences in TFs at the protein level and subsequent changes in specific groups of transcripts regulated by those TFs.

Interestingly, the analysis of the correlation of expression with HAX1, performed for TCGA database in four neoplasms (AML, DLBCL, breast cancer, cervical cancer) identified enrichment in the same biological processes as detected for *HAX1* KO vs. WT in HL-60 cell line, indicating that these results are not cell-line or neoplasm-specific and further corroborating these findings.

Observed changes in expression linked to ribosome assembly and translation are not huge, but persistent, pertain to relatively large group of transcripts and seem not to be cell type or neoplasm-specific. Relatively weak binding and small, but reliable changes in expression suggest the regulation via small, additive effects. It is an open question if these effects can manifest more robustly in non-quiescent cells, subjected to some kind of stress. This conjecture is supported by the reported changes in localization of HAX1 upon stress, including nucleocytoplasmic shuttling (Grzybowska et al. 2013), which could be linked to ribosome biogenesis and RNA binding in the nucleus/nucleolus. Moreover, the observed abundance of rRNA as a potential target in CRAC analysis suggests the possibility of more direct involvement in ribosome biogenesis. The suggested binding site (Figure S3) maps within rRNA expansion segments, for which the function in ribosome biogenesis was proposed (Ramesh and Woolford 2016). Thus, HAX1 involvement in ribosome assembly might encompass not only regulation of stability of mRNAs encoding ribosomal proteins and assembly factors, but also a direct interaction with ribosomal RNA. Interestingly, the possibility of the simultaneous regulation of translation by direct ribosome binding and controlling mRNA stability was described for FMRP protein (Chen et al. 2014), already mentioned here for similar mode of binding and possible recruitment of transcripts to P-bodies.

To test physiological consequences of *HAX1* KO on ribosome status we performed ribosome sedimentation profiling, demonstrating a difference in 40S:monosome ratio for HAX1-deficient cells, and, to less extent, in 40S+60S:monosome ratio. Shift towards free subunits in *HAX1* KO may indicate less efficient monosome assembly, resulting from lower expression of ribosomal assembly factors and ribosomal components.

In conclusion, we provide evidence for HAX1 involvement in ribosome biogenesis and translation, which represents a new finding. Previously, it was reported that HAX1 has an interaction with PELO, a protein involved in ribosomal rescue during ribosome stalling, but no mechanism for HAX1 involvement was proposed and no physiological effect was observed (Burnicka-Turek et al. 2010). Recently, You et al. (You et al. 2022) demonstrated, among other things, that HAX1 levels correlate with ribosome formation and that HAX1 promotes translation of the transcript encoding integrin subunit beta 6 in endothelial cells, which partially tallies with the findings presented here.

Thus, presented results suggest a possibility that HAX1 binds to the CDS of the non-translated transcripts, protecting from degradation and that the main mRNA targets subjected to this regulation include transcripts involved in ribosome biogenesis.

Changes in ribosomal status and the efficiency of translation may affect proliferation, which in turn may contribute (in opposite directions) to neutropenia and/or cancer. Indeed, HAX1 status has been already shown to affect proliferation (Wu et al. 2017) and we also observed this effect in our cell lines (data not shown). Further research should elucidate the exact role of HAX1 in maintaining translation efficiency, but the results presented here provide a starting point to explore these new and unanticipated possibilities.

## AVAILABILITY

HAX1 targets genomic position available at UCSC Genome Browser (http://genome-euro.ucsc.edu/s/Grzegorz%20Kudla/Ewelina_HAX_MCPIP)

## ACCESSION NUMBERS

NGS data have been deposited with the Gene Expression Omnibus (GEO) functional genomics data repository, accession numbers: GSE189611, GSE189609.

## SUPPLEMENTARY DATA

Supplementary Figures: S1-S5

Supplementary files: S1-S5

## ACKNOWLEDGEMENT

The authors are grateful to Alicja Trebinska-Stryjewska and Antek Laczkowski for helpful comments.

