## Supplemental data file for "The RNA-binding landscape of HAX1 protein indicates its involvement in ribosome biogenesis and translation"

SUPPLEMENTARY DATA

Figures:

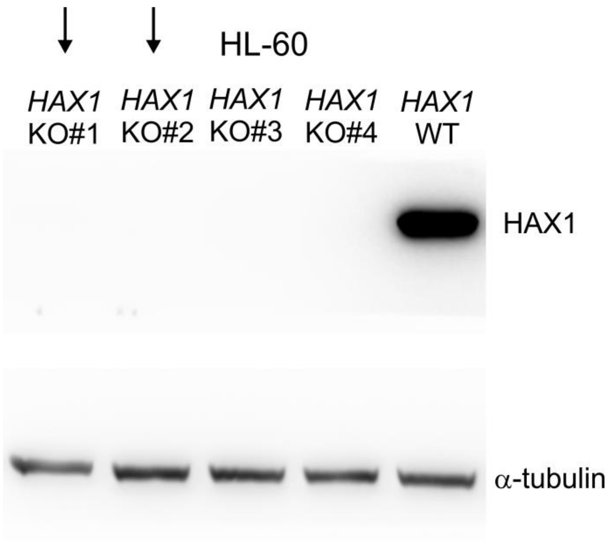

Figure S1. Western blot with four established *HAX1* KO cell lines (#1-4) and the wild type (HL-60 cell line). α-tubulin represents the reference. HAX1 KO cell lines marked with arrows were used in RNA-seq and sedimentation experiments.

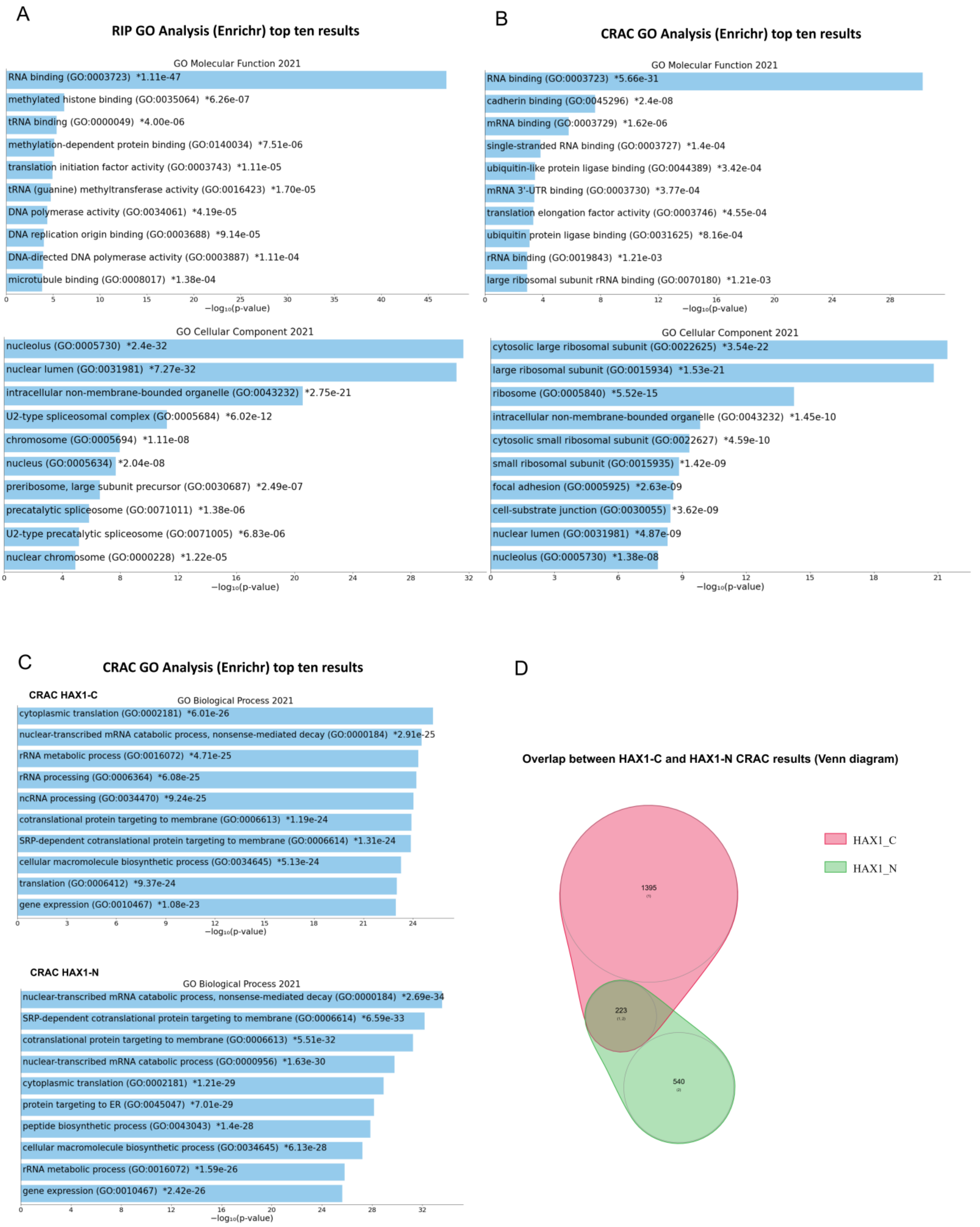

Figure S2. Functional annotation enrichment of HAX1 target genes in RIP-seq and CRAC analyses.

A. Enriched Gene Ontology terms (Enrichr Ontologies: Molecular Function, Cellular Component) in RIP-seq analysis presented in bar plot, the length of each bar corresponds to statistical significance of the enrichment (-log10 from adjusted p-value). Top ten terms are presented. B. Enriched Gene Ontology terms (Enrichr Ontologies: Molecular Function, Cellular Component) in CRAC analysis (joint HAX1-C and HAX1-N results) presented in bar plot, the length of each bar corresponds to statistical significance of the enrichment (-log10 from adjusted p-value). Top ten terms are presented. C.

Separate analysis of CRAC experiments with the tag on C-terminus (HAX1-C) and N-terminus (HAX1-N): Gene Ontology analysis (Biological Process 2021) for both experiments (Enrichr, ten top terms, plot created by Appyter) D. Venn diagram demonstrating the overlap between the results of HAX1-C and HAX1-N experiments.

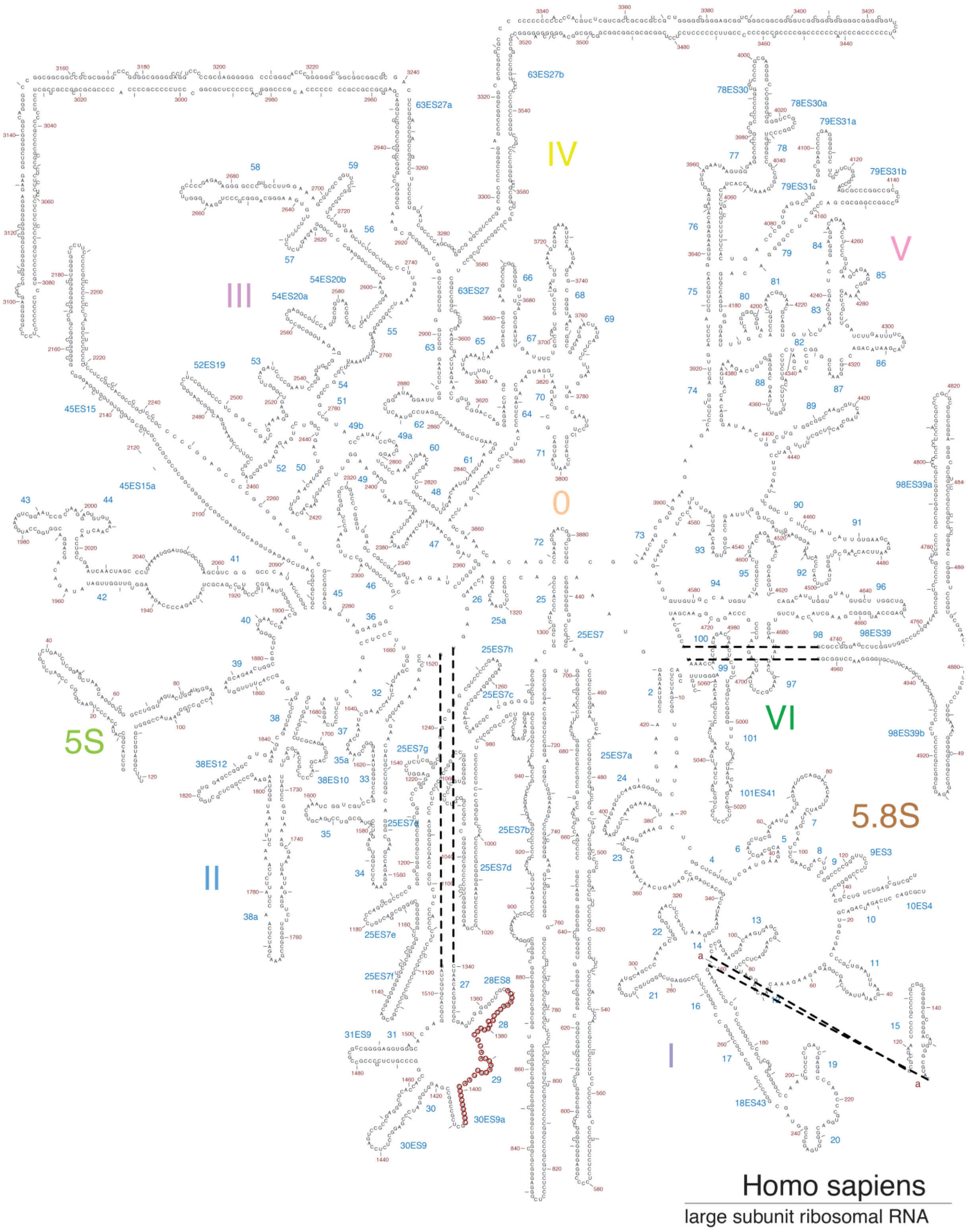

Figure S3. Large ribosomal subunit rRNA with marked region of potential interaction site with HAX1 (nucleotides marked by red circles).

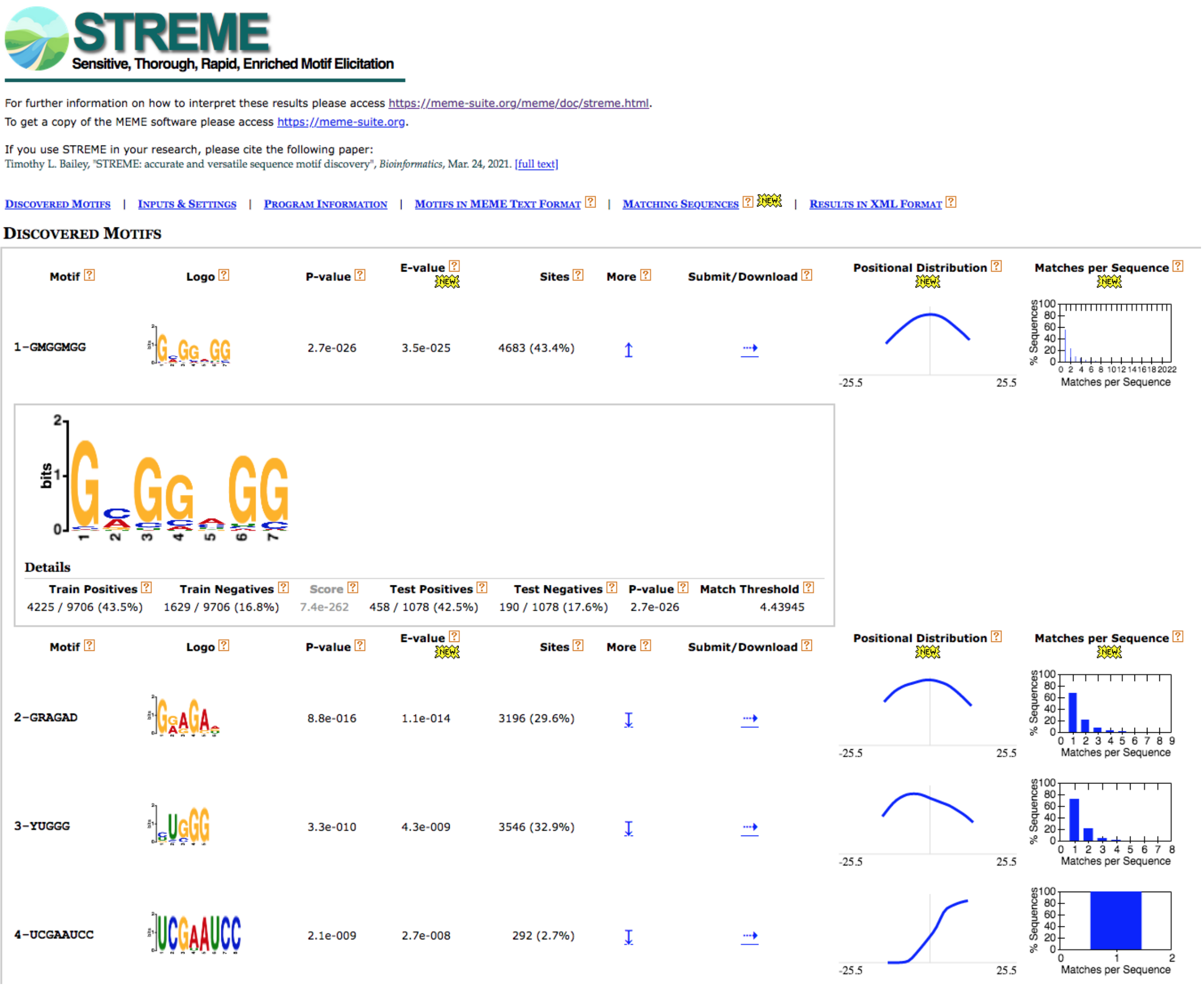

Figure S4. STREME analysis of combined CRAC data for HAX1-binding motif in mRNA targets.

A

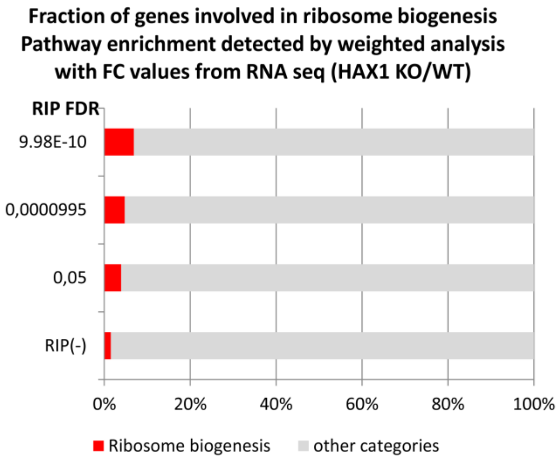

B

| TF | RIP FC | RIP adjusted p-value | Enrichr Combined Score (ENCODE) |
|---|---|---|---|
| MYC | 1,204498 | 3,54E-09 | 302,66 |
| TAF1 | 1,482814 | 1,11E-15 | 109,54 |
| YY1 | 2,049345 | 3,7E-106 | 86,95 |
| E2F6 | 1,431905 | 2,19E-12 | 86,83 |
| BRCA1 | 1,200453 | 1,15E-07 | 62,52 |
| USF1 | 4,655167 | 7,3E-104 | 16,48 |
| MYCBP | 3,285746 | 3,1E-113 | |
| MYCBP2 | 1,287111 | 1,29E-08 | |

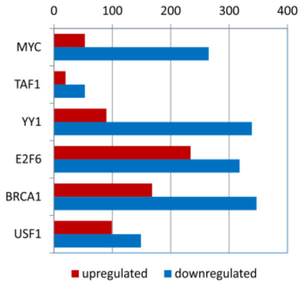

Figure S5. Analysis of different expression profile resulting from *HAX1* knockout in HL-60 cell line.

A. Fraction of transcripts involved in ribosome biogenesis increases with decreasing FDR for RIP target subset. Pathway enrichment calculated in String11 weighted analysis with FC values from RNA-seq (RNA-seq_*HAX1*_KO/RNA-seq_WT). B. Analysis of transcription factors possibly affected by *HAX1* KO. Upper panel: a list of the most probable transcription factors identified by Enrichr as regulating a subset of genes with significantly changed expression in *HAX1* KO simultaneously identified as RIP targets. The two MYC-binding proteins are included as high-scoring in RIP and with a potential of influencing the performance of MYC . Lower panel: a bar plot showing the number of upregulated and downregulated genes in *HAX1* KO for the specific TFs.

Supplementary files description (Excel files):

File S1. RIP-seq results; experimental data and analysis.

Sheet1 (all): results for all genes (pulldown: HAX1/IgG)

Sheet2 (positive): only positive targets with FC and FDR

Sheet3 (RNA classes): RNA biotypes detected in RIP

Sheet4 (Enrichr_BP): enrichments in Biological Process for RIP targets (Enrichr)

Sheet5 (Enrichr_MF): enrichments in Molecular Function

Sheet6 (Enrichr_CC): enrichments in Cellular Component

Sheet7 (Panther GO BP): enrichments in Slim Biological Process (Gene Ontology Panther)

File S2. CRAC results; experimental data and analysis.

Sheet1 (mRNA_counts_hOH7 FC): results for all genes (HAX1 overexpressing cell line/control cell line)

Sheet2 (Gene_Symbol_N_C_overlap): targets overlapping in N and C datasets

Sheet3 (Enrichr_BP): enrichments in Biological Process for pooled CRAC targets (Enrichr)

Sheet4 (Enrichr_MF): enrichments in Molecular Function

Sheet5 (Enrichr_CC): enrichments in Cellular Component

Sheet6 (Panther GO BP): enrichments in Slim Biological Process (Gene Ontology Panther)

Sheet7 (Enrichr_BP_HAX_C): enrichments in Biological Process for HAX1_C CRAC targets (Enrichr)

Sheet8 (Enrichr_BP_HAX_N): enrichments in Biological Process for HAX1_N CRAC targets (Enrichr)

Sheet9 (RNA classes): RNA biotypes detected in CRAC

Sheet10 (Genomic position): genomic position of pooled CRAC targets

File S3. Overlap of RIP and CRAC results

Sheet1 (Gene name): Lists of gene names representing significant results obtained in RIP and CRAC and the overlapping part

Sheet2 (Enrichr_BP): enrichments in Biological Process for the overlapping targets (Enrichr)

Sheet3 (Enrichr_MF): enrichments in Molecular Function

Sheet4 (Enrichr_CC): enrichments in Cellular Component

Sheet6 (Panther GO BP): enrichments in Biological Process Complete (Gene Ontology Panther)

File S4. RNA-seq (HAX1 KO/WT) results.

Sheet1 (all): results for all genes, FC calculated as a combined value for HAX1 KO#1 and HAX1 KO#2

Sheet2 (downregulated): only significantly downregulated genes

Sheet3 (upregulated): only significantly upregulated genes

Sheet4 (GO_Biological Process_String11): String11 weighted analysis of the results with FC(RNA-seq HAX1 KO/RNA-seq WT) as values.

File S5. Correlation of expression of HAX1 with other genes in different neoplasm (cBioPortal analysis)

Sheet1 (InvBC_genes): Invasive Breast Cancer (TCGA, PanCancer Atlas), gene names and correlation assessment (Spearman coefficient, p-value, q-value)

Sheet2 (InvBC_BP): Invasive Breast Cancer, String 11 GO weighted analysis with Spearman coefficient as values, Biological Process terms

Sheet3 (Cervical_carcinoma_genes): Cervical Carcinoma (TCGA, PanCancer Atlas), gene names and correlation assessment (Spearman coefficient, p-value, q-value)

Sheet4 (Cervical Carcinoma_BP): Cervical Carcinoma, String 11 GO weighted analysis with Spearman coefficient as values, Biological Process terms

Sheet5 (AML_genes): Acute Myeloid Leukemia (TCGA, PanCancer Atlas), gene names and correlation assessment (Spearman coefficient, p-value, q-value)

Sheet6 (AML_BP): Acute Myeloid Leukemia, String 11 GO weighted analysis with Spearman coefficient as values, Biological Process terms

Sheet7 (DLBCL_genes): Diffuse Large B Cell Lymphoma (TCGA, PanCancer Atlas), gene names and correlation assessment (Spearman coefficient, p-value, q-value)

Sheet8 (DLBCL_BP): Diffuse Large B Cell Lymphoma, String 11 GO weighted analysis with Spearman coefficient as values, Biological Process terms
